# Microtubules in Arabidopsis pollen tubes are oriented away from the tube apex and are actin-independent at the cortex

**DOI:** 10.64898/2026.01.21.700958

**Authors:** Joshua H. Coomey, Eden R. Gallup, Ram Dixit

**Author notes:** denotes equal contribution (Joshua H. Coomey, Eden R. Gallup). The author responsible for distribution of materials integral to the findings presented in this article in accordance with the policy described in the Instructions for Authors (https://academic.oup.com/plphys/pages/General-Instructions) is Ram Dixit.

## Abstract

Pollen tubes are dynamic tip-growing cells that deliver sperm nuclei to female gametes in flowering plants, allowing for sexual reproduction and seed formation. Actin and microtubule cytoskeletons both play important roles in directional pollen tube growth and guidance. While actin dynamics are well-studied in pollen tubes, the role of microtubules and the interactions between these two cytoskeletal filaments are less well understood. To address this knowledge gap, we imaged growing *Arabidopsis thaliana* pollen tubes co-expressing fluorescently-labeled tubulin and actin markers and observed partial co-localization of actin and microtubule filaments. We found that treatment with microtubule disrupting drugs did not affect the actin cytoskeleton. In contrast, when actin filaments were depolymerized, microtubules in the medial region of pollen tubes were disrupted, while microtubules at the cell cortex remained intact. Thus, the microtubule cytoskeleton in *A. thaliana* pollen tubes relies on the actin cytoskeleton in a spatially dependent manner. Furthermore, we utilized native expression of the microtubule plus-end binding protein EB1b to track microtubule orientation in growing pollen tubes. We found the microtubule array to be largely parallel, with plus ends growing away from the tube apex. Together, these findings offer new insights into the dynamics and organization of microtubules in growing pollen tubes and the interactions between actin filaments and microtubules.

## INTRODUCTION

In flowering plants, pollen tubes carry the male gamete to the ovule located deep within the carpel tissue. Thus, pollen function is essential for successful fertilization which underlies most of our fruit and grain crops. Beyond agricultural applications, pollen is also a unique and important system for understanding how cells respond to mechanical demands and facilitate tip growth. Pollen undergoes mechanically intensive processes in order to fertilize the female gamete; it first undergoes dehydration to allow for dispersal from the anther, and upon landing on a compatible stigma, it rapidly rehydrates and becomes metabolically active (Jackson, 1989). Hydrated pollen germinates a tip-growing pollen tube, which penetrates through the style to reach the ovary, where it then bursts to release the sperm cells in the ovule (Krichevsky et al., 2007). The pollen tube is the fastest-growing cell in the plant kingdom, reaching speeds of up to 2.8 µm/s in maize (Selinski and Scheibe, 2014).

Tip growth requires polarization of the cell toward the growing tip and targeted secretion to drive tip elongation (Cai et al., 2015). The cytoskeleton plays a crucial role in both processes. Actin filaments are necessary for pollen tube germination and elongation. When actin disrupting drugs are applied, pollen tube growth is immediately arrested (Vidali et al., 2001; Xu and Huang, 2020). Additionally, the actin cytoskeleton is the main driver of cytoplasmic streaming, the process by which organelles and vesicles containing cell wall material are transported by actin-dependent myosin motor proteins to the tube apex to facilitate tip growth, while excess plasma membrane is internalized and recycled (Hepler et al., 2001; Cheung et al., 2008).

The function of microtubules in the pollen tube is less well understood. Treatment with microtubule depolymerizing agents has little effect on pollen tube growth rate, indicating a nonessential role in this process (Heslop-Harrison et al., 1988; Gossot and Geitmann, 2007). Studies have suggested that microtubules play a role in callose deposition (Cai et al., 2011; Parrotta et al., 2022), pollen tube growth directionality *in vitro* (Gossot and Geitmann, 2007), and in transportation of the sperm cells through the pollen tube (Schattner et al., 2020; Wang et al., 2024). One theory proposes that actin and microtubules may work together, with multiple actin and microtubule motor proteins bound to an organelle at the same time, resulting in organelles that move at different speeds depending on the type and direction of the motor proteins to which they are bound (Cai and Cresti, 2006; Cai et al., 2015).

Since motor proteins travel unidirectionally along cytoskeletal filaments, the array orientation is a key determinant of directional delivery of cargo in cells. In root hairs, microtubules are oriented with their plus ends facing away from the nucleus, both towards the root hair tip and towards the root, and mediate aspects of polarized growth (Bibikova et al., 1999; Ambrose and Wasteneys, 2014). In moss protonema, microtubules are largely oriented with plus ends facing towards the cell apex, forming a focal point in the apex that is associated with tip growth (Doonan et al., 1988; Hiwatashi et al., 2014; Wu and Bezanilla, 2018). However, the orientation of microtubules in pollen tubes remains unclear.

By generating new dual-marker lines that fluorescently label both the actin and microtubule cytoskeletons in *A. thaliana* pollen, we have determined that microtubules and actin filaments partially colocalize in growing pollen tubes. We pharmacologically disrupted both cytoskeletal filaments and found that while actin filaments do not depend on the presence of microtubules, disruption of actin leads to differential degradation of the microtubule cytoskeleton based on location in the pollen tube. Additionally, we used a natively expressed EB1b-GFP reporter to track microtubule plus-end dynamics and found the pollen tube microtubule array to be largely oriented away from the tube apex, in contrast to other tip growing cells. Together, these data reveal novel features of microtubule organization in pollen tubes.

## MATERIALS AND METHODS

### Generation of transgenic lines

*A. thaliana* plants expressing *Lat52::mRuby2-TUB6* were generated using multisite-Gateway recombination in pB7WGLat52 vector and introduced into Col-0 (wildtype) plants using *Agrobacterium tumefaciens*-mediated transformation. Similarly, *Lat52:LifeAct-mEGFP* lines were generated and the two populations were crossed to produce plants expressing both fluorescent markers. The *Lat52::mRuby2-TUB6* construct was also transformed into the *proEB1b::EB1b-mEGFP* marker line (Zhang et al., 2013) to enable visualization of both microtubules and their growing plus ends. All plant lines were grown under long-day conditions (16 h light/8 h dark), in 175 µmol light, 45% relative humidity, 21°C. For each line, multiple independent transformants were selected and they all showed normal seed set, pollen germination, pollen growth rate, and overall plant morphology.

### Pollen germination and imaging

Plants were grown until flowering (approximately 5 weeks) in long-day conditions and pollen was germinated as described previously (Coomey and Haswell, 2023). Approximately 40 flowers were picked using tweezers between 9:00am and 11:30am and placed in a 1.7 mL Eppendorf tube with 750 µL standard liquid pollen growth medium (LPGM; 0.49 mM H_3_BO_3_, 2 mM Ca(NO_3_)_2_, 2 mM CaCl_2_, 1 mM KCl, 1 mM MgSO_4_, and 18% (w/v) sucrose with pH 7.05-7.10 adjusted with KOH). The mixture was vortexed for 2 min and centrifuged at 10,000 x g for 4 min before supernatant and flower material were removed and pollen was resuspended in 15 µL fresh LPGM. Pollen was transferred to a glass microwell dish (MatTek 35mm dish, No. 1.5 coverslip, 14mm glass diameter, uncoated) pre-treated with polyethyleneimine (PEI, to adhere pollen to glass) using a cut pipet tip to accommodate the pollen size. Pollen was spread evenly on the dish and allowed to dry and adhere to PEI for 1-2 minutes, until it acquired a grainy appearance. 200 µL LPGM was added carefully to the edge of the dish to avoid disturbing the pollen. Dishes were placed in a large glass Petri dish lined with a wet Kimwipe to maintain humidity and incubated at 22°C for 3-4 hours in the dark.

### Imaging and colocalization analysis

Pollen was imaged on an Olympus FluoView 3000 laser scanning confocal microscope with UPLSAPO 60XW NA1.2 water-immersion objective. Samples were excited with 488 nm laser to visualize LifeAct-mEGFP and EB1b-mEGFP, and with 561 nm laser to visualize mRuby2-TUB6. Emitted fluorescence was captured at 500-540 nm (mEGFP) and 585-673 nm (mRuby2) using sequential scanning. Images were deconvolved using the Olympus cellSens constrained iterative deconvolution.

Signal colocalization was analyzed using the BIOP JACoP plugin for ImageJ. A freehand ROI was drawn around the pollen tube, accommodating the entire Z stack, and images were auto-thresholded using the IJ_Isodata thresholding method. Pearson Correlation Coefficient (PCC) and Manders’ Correlation Coefficients (MCC) were generated and used to interpret colocalization, with PCC showing the strength of linear correlation between actin and microtubule signal (on a scale -1 to 1, with 1 representing complete colocalization and -1 representing complete exclusion), and MCC M1 and M2 representing the percentage of actin signal coinciding with microtubule signal, and the percentage of microtubule signal overlapping with actin signal, respectively.

### Cytoskeleton depolymerization experiments

Pollen co-expressing *Lat52::mRuby2-TUB6* and *Lat52::LifeAct-mEGFP* was germinated for 3-4 hours and imaged by confocal laser scanning microscope as described above. Before imaging, LPGM in the glass dish was carefully pipetted off and replaced with 100 µL of LPGM with 1.5 µM oryzalin or 30 nM latrunculin B, or an equivalent amount of DMSO (0.15% and 0.30%, respectively). Oryzalin concentration was determined by testing a range of concentrations from 1.5 µM to 100 µM and selecting the minimum concentration that resulted in microtubule depolymerization. Latrunculin B concentration was determined through a review of similar experiments in *A. thaliana* and other species (Cheung et al., 2008; Zhu et al., 2017). Pollen was incubated with the drugs for at least 10 min, and imaging was completed within 45 min of treatment.

To analyze the effects of these treatments, actin and microtubule cytoskeletons in each pollen tube image were classified as “none” (no filaments visible), “minimal” (1-5 filaments visible), or “typical” (6+ filaments visible, actin/microtubules appear unaffected). To further assess the location of microtubules after treatment, the microtubule cytoskeleton in each pollen tube image was classified as “medial,” “cortical,” “medial and cortical,” and “none” based on the distance of tubulin signal from the periphery of the pollen tube, assessed visually by examining the image z-stack.

## RESULTS

### Microtubules and actin filaments partially colocalize in growing pollen tubes

To investigate the spatial distribution of actin filaments and microtubules in *A. thaliana* pollen tubes, we utilized dual-reporter lines co-expressing fluorescently-labeled actin and tubulin under the pollen-specific Lat52 promoter (*Lat52::LifeAct-mEGFP* and *Lat52::mRuby2-TUB6*). Confocal laser scanning microscopy of live pollen tubes revealed characteristic organization of the actin and microtubule cytoskeletons. The actin cytoskeleton consisted of a “fringe” of short, dynamic actin filaments at the tube apex, as well as long actin cables down the shank of the pollen tube (Fig. 1A). This pattern is consistent with previous literature describing the organization of actin filaments in tobacco and lily pollen tubes (Gossot and Geitmann, 2007; Cheung et al., 2008). The mRuby2-TUB6 signal showed short, sparse filaments near the tube apex, with longer filaments visible in the shank, largely parallel to the axis of growth (Fig. 1B).

**Figure 1.**
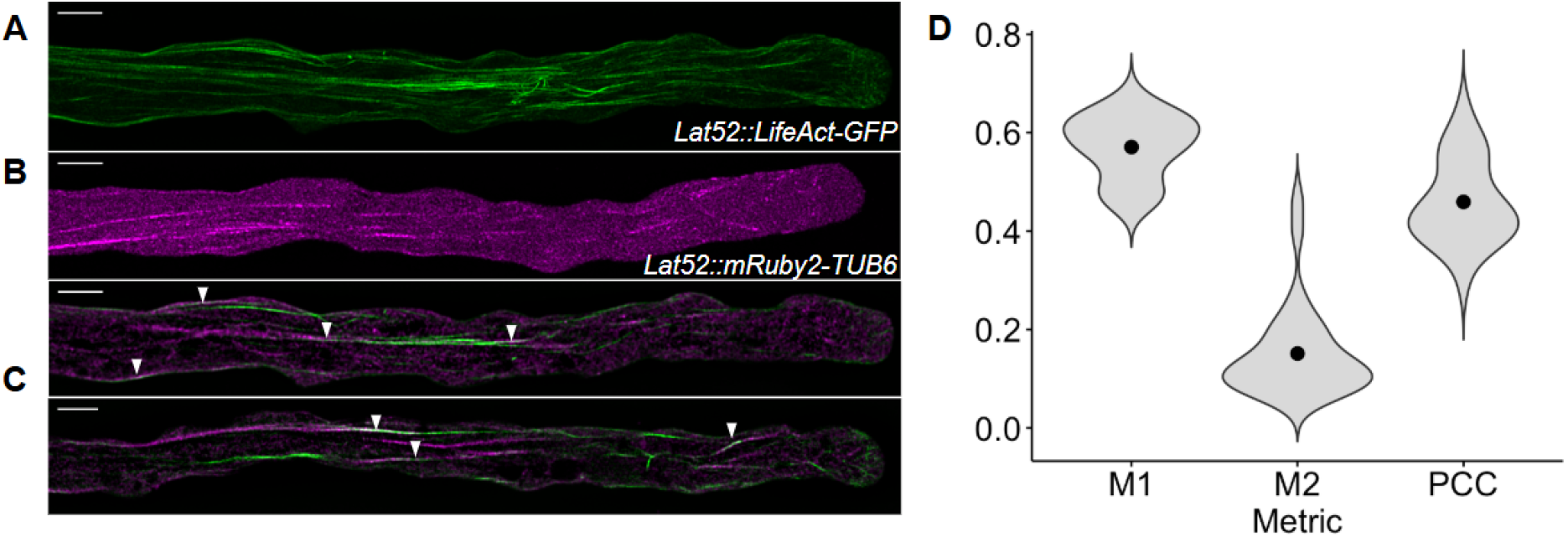
Microtubules and actin filaments display distinct organization and partial co-localization in the *A. thaliana* pollen tube. **A)** The actin cytoskeleton in a growing pollen tube shows a fringe near the tube apex and long cables down the shank. Image is a maximum Z projection. **B)** Microtubules in a growing pollen tube exist as long parallel filaments in the medial shank and as sparse short fragments near the tube apex. Image is a maximum Z projection. **C)** Two images of pollen tubes expressing both actin and tubulin markers. Both images are single Z-slices from medial sections of the tubes; the top image is from the same pollen tube shown in (A). Examples of colocalization events are marked with arrows. **D)** Quantification of actin and tubulin signal colocalization using Manders (M1 = percentage of actin signal overlapping with tubulin signal. M2 = percentage of tubulin signal overlapping with actin signal) and Pearson correlation coefficients (n = 30 pollen tubes). Scale bars = 5 μm.

Imaging also showed partial colocalization of the actin filaments and microtubules. Segments of filamentous LifeAct-mEGFP and mRuby2-TUB6 signal frequently overlapped spatially in the pollen tube, although there were also many examples of non-overlapping filaments for both reporters (Fig. 1C). We quantified the colocalization between the actin and microtubule markers using two metrics: the Pearson correlation coefficient (PCC) which measures the covariance of the signal intensities in each pixel of the two images, and the Manders’ colocalization coefficients (M1 and M2) which measure the reciprocal signal overlap between the two images (Fig. 1D). The PCC of the pollen tube images averaged 0.46 (n = 30), which indicates a moderate association consistent with the partial colocalization visible in the images. The Manders’ colocalization coefficients M1 and M2 measured the percentage of actin signal overlapping tubulin signal, and the percentage of tubulin signal overlapping actin signal, respectively. They averaged to M1 = 0.57 and M2 = 0.15 (n = 30). The low M2 value indicates that fewer microtubules are associated with actin filaments. Alternatively, it is possible that the relatively high level of cytoplasmic signal in the microtubule marker line produces a deflated estimate of the amount of microtubule signal that coincides with actin filaments.

### Depolymerization of the actin cytoskeleton disrupts microtubules in a spatially-dependent manner

Previous research in *Papaver rhoeas* pollen suggests that the actin cytoskeleton does not rely on the presence of microtubules, while pharmacologically depolymerizing actin filaments results in complete degradation of microtubules (Gossot and Geitmann, 2007). To test whether these effects are conserved in *A. thaliana* pollen tubes, we first treated our dual-reporter pollen tubes with 1.5 µM oryzalin (Fig. 2A). As expected, oryzalin treatment resulted in complete depolymerization of microtubules, and no discernible change in the actin cytoskeleton compared to DMSO control (Fig. 2C).

**Figure 2.**
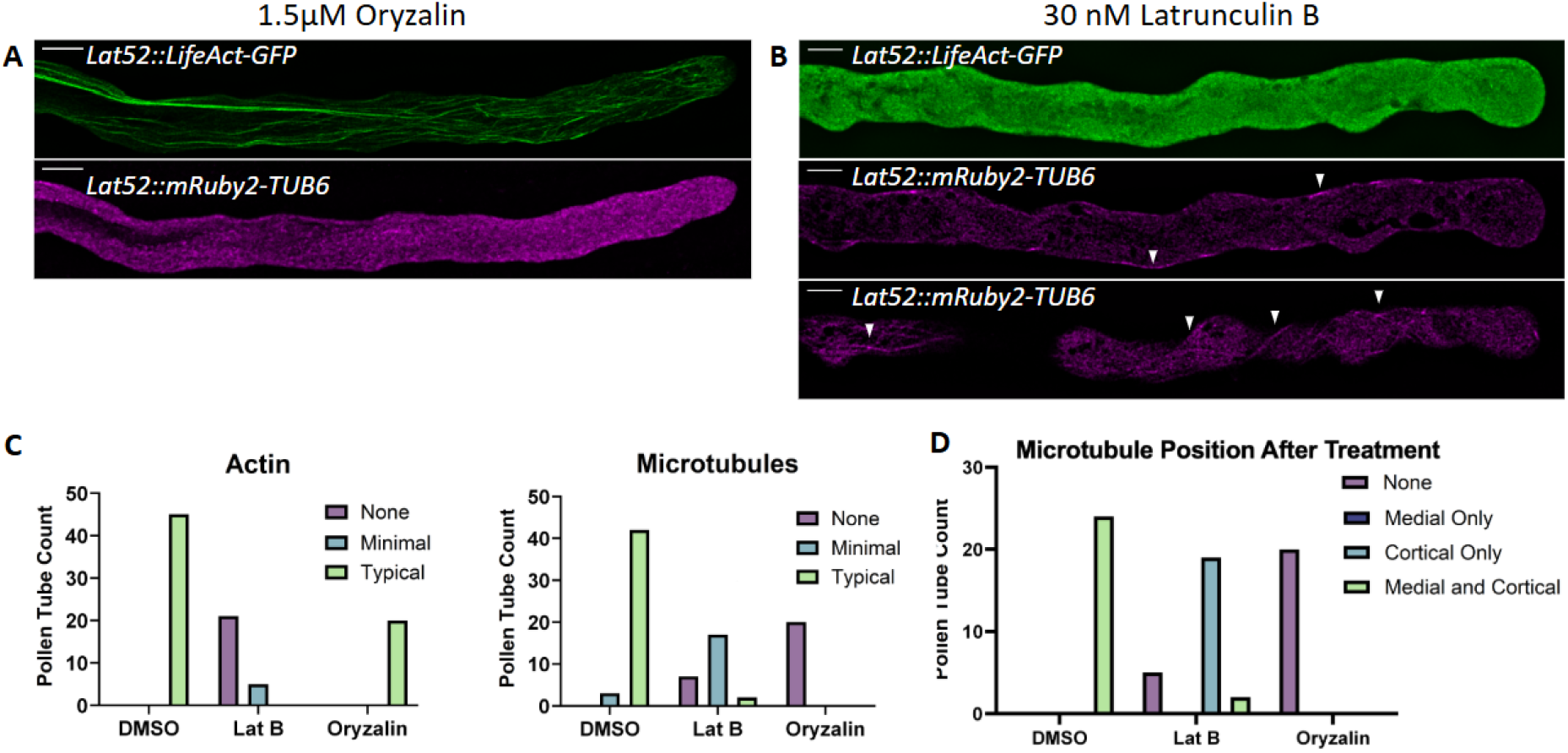
Medial microtubules are actin-dependent while cortical microtubules are actin-independent. **A)** Representative maximum Z projection images of actin (top) and tubulin (bottom) marker after treatment with 1.5 μM oryzalin. Microtubules are completely depolymerized, while actin filaments are essentially unaffected. **B)** Representative images of pollen tubes after treatment with 30 nM latrunculin B. Top image is a maximum Z projection showing complete depolymerization of actin filaments. Middle image is a single medial Z slice, and the bottom image is a single cortical Z-slice showing disruption of filamentous tubulin signal in the medial portion of the pollen tube, and microtubules persisting in the cortical region (marked with arrows), respectively. **C)** Effect of treatment with latrunculin B (n = 26), oryzalin (n = 20), or DMSO (n = 24, n = 21) on actin filaments and microtubules. P < 0.0001 by Fisher’s Exact Test. **D)** Position of the microtubules remaining after treatment with latrunculin B (n = 26), oryzalin (n = 20) and DMSO (n = 24). P < 0.0001 by Fisher’s Exact Test. Scale bars = 5 μm.

We then treated dual-reporter pollen tubes with 30 nM latrunculin B (Fig. 2B). While actin was largely depolymerized after treatment, microtubules were only partially disrupted (Fig. 2C). A closer inspection of the remaining microtubules revealed that they were overwhelmingly located in the cortical region of the pollen tube shank, while microtubules in the medial region of the pollen tube were almost completely depolymerized (Fig 2D). This finding reveals that medial and cortical pollen microtubules are differentially sensitive to the loss of actin filaments in *A. thaliana*.

### Microtubules are oriented predominantly with plus ends growing away from the tube apex

We next wanted to investigate the orientation of the microtubule array in pollen tubes. We used the plus-end tracking protein, EB1b, expressed under its native promoter (*proEB1b::EB1b-mEGFP*) to track the direction of growing microtubules in the apical, mid-shank, and distal shank regions of growing pollen tubes (Fig. 3A). We imaged growing pollen tubes every 4 sec for 2 min total, collecting short Z-stacks from the top plane of the cortex to the medial region.

**Figure 3.**
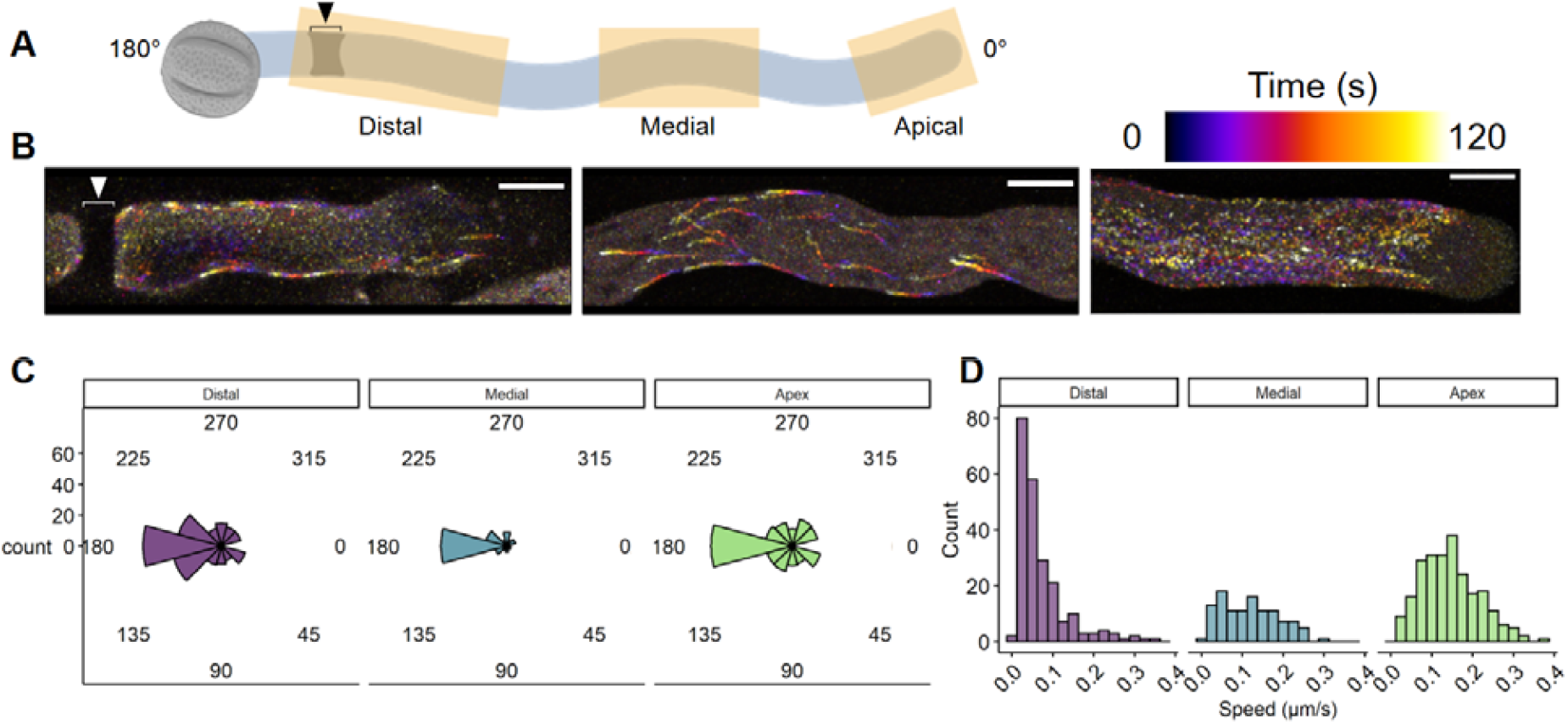
Microtubules are oriented away from the pollen tube apex. **A)** Schematic of a pollen tube showing direction of growth (0°) away from the grain (180°). Yellow highlights indicate imaged regions. Black arrow and bracket indicate callose plug. **B)** Representative color-coded time series of EB1b signal in imaged regions over a 120s duration. Color scale indicates time point. White arrow and bracket indicate callose plug. Scale bar = 5 μm. **C)** Quantification of EB1b trajectories from the indicated regions of 8 individual pollen tubes: distal (n = 225), medial (n = 112), and apical (n = 238). **(D)** Distribution of EB1b particle speed from the indicated regions.

EB1b particle trajectory and speed were then quantified using the Particle Analysis plugin in ImageJ. In the apical region, there were numerous EB1b particles with mixed trajectories (Fig. 3B, Supplemental Movie 1). In the mid-shank and distal shank regions, EB1b particles had more uniform trajectories (Fig. 3B, Supplemental Movies 2 and 3). Across all regions, we observed that the majority of EB1b particles were oriented away from the tube apex (Fig. 3C). EB1b average particle speed was somewhat slower in distal regions (0.07 ± 0.06 µm/s), with medial and apical regions showing similar speeds (0.11 ± 0.07 µm/s and 0.14 ± 0.07 µm/s respectively), all consistent with speeds reported in other systems (Fig. 3D) (Liang et al., 2009; Harris et al., 2016). These data show that the microtubule array in *A. thaliana* pollen tubes consists largely of parallel microtubules, oriented with plus ends growing away from the apex.

## DISCUSSION

Using a new *A. thaliana* actin and tubulin dual-reporter pollen line, we have determined that the actin and microtubule cytoskeletons partially colocalize in live pollen tubes, consistent with previous findings using electron microscopy (Lancelle and Hepler, 1991). Pharmacological depolymerization of the two cytoskeletal components demonstrated that the arrangement of the actin cytoskeleton is independent of microtubules as reported previously (Gossot and Geitmann, 2007), whereas a subpopulation of microtubules is dependent on the presence of actin filaments. Specifically, microtubules in the medial region of the pollen tube are disrupted in response to actin depolymerization, while microtubules at the cell cortex remain largely unaffected. These data suggest that attachment of microtubules to the cell cortex makes them resistant to actin disrupting drugs. Further investigation is needed to determine whether the interactions between microtubules and actin filaments occur through direct or indirect mechanisms. Potential factors that might mediate microtubule-actin crosstalk in pollen tubes are KCH kinesins (Schattner et al., 2020) and microtubule-associated proteins such as SB401 (Huang et al., 2007).

Lipid composition has emerged as another possible mechanism to coordinate the activities of the actin and microtubules cytoskeletons. Recent work has found that detergent-insoluble membrane microdomains from tobacco pollen contain both tubulin and actin (Onelli et al., 2025). Reducing the sterol content decreases the number of microtubules, alters microtubule patterning, and inhibits microtubule regrowth following depolymerization (Onelli et al., 2025). Thus, variation in plasma membrane and endomembrane lipid composition might contribute to the medial actin-dependent and cortical actin-independent microtubule populations we observed. The nature of these spatially distinct microtubule populations and their relevance to pollen tube growth remains to be explored.

Our work used native EB1b expression to study microtubule polarity in pollen tubes. Importantly, endogenous expression of EB1b labeled growing microtubule plus ends in the expected comet-like fashion instead of labeling large segments of the pollen microtubules as observed upon overexpression of EB1b (Chan et al., 2003; Wang et al., 2024). Our finding that the pollen tube microtubules are largely oriented with plus ends growing away from the cell apex sets this system apart from other tip-growing cell types. In root hairs and moss protonema, the microtubules are largely facing towards the cell apex, with plus-end directed motor proteins delivering cargo to the growing tip (Ambrose and Wasteneys, 2014; Hiwatashi et al., 2014; Wu and Bezanilla, 2018). In fission yeast, the interaction of microtubule plus ends is involved with defining sites for polar cell growth (Minc et al., 2009). The largely proximal-facing microtubule orientation we observe in *A. thaliana* pollen tubes suggests a shift in polar growth mechanics relative to other tip growing systems and is consistent with the well-documented continuation of tip growth when microtubules are chemically depolymerized. Our data are also consistent with the finding that two members of the kinesin-14 family, which are expected to move towards the minus-end of microtubules, contribute to the tipward transport of the male germ unit (Yan et al., 2025). A better understanding of microtubule orientation and actin-microtubule interactions lays the foundation to investigate the factors regulating these processes and how they contribute to the mechanics of pollen tube growth.

## Acknowledgments

This work was supported by the National Institute of General Medical Sciences of the National Institutes of Health under award number R35GM139552 (R.D.).

## Author Contributions

J.C., E.G., and R.D. conceived of the study. J.C. and E.G. carried out the experiments and performed the analysis. J.C., E.G., and R.D. interpreted the data and wrote the manuscript.

## SUPPLEMENTAL INFORMATION

**Supplemental Movie 1.**
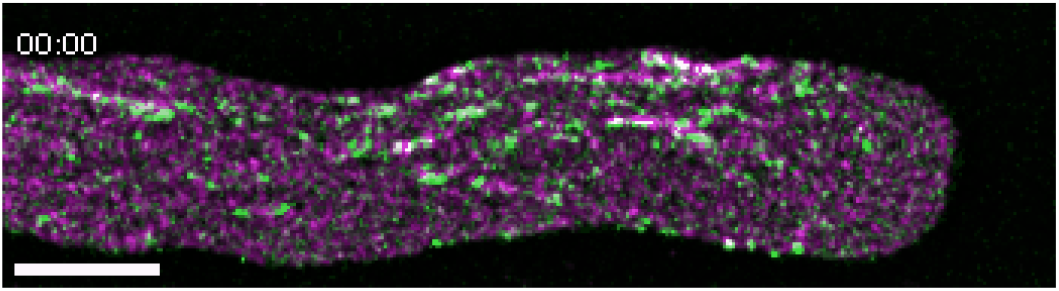
Timelapse of EB1b-mEGFP and mRuby2-TUB6 signal in the pollen tube apex. Images were captured every 10 seconds. Scale bar = 5 μm.

**Supplemental Movie 2.**
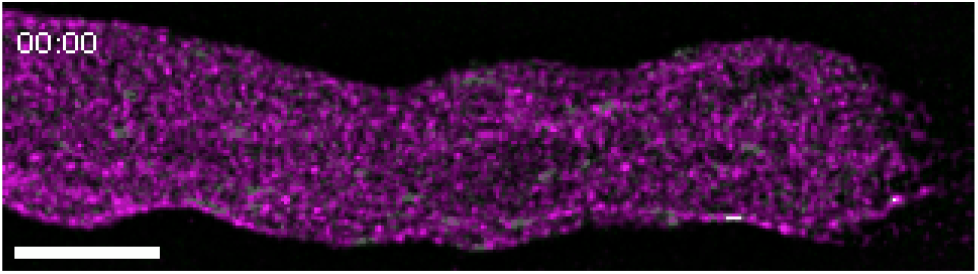
Timelapse of EB1b-mEGFP and mRuby2-TUB6 signal in the pollen tube mid-shank region. Images were captured every 10 seconds. Scale bar = 5 μm.

**Supplemental Movie 3.**
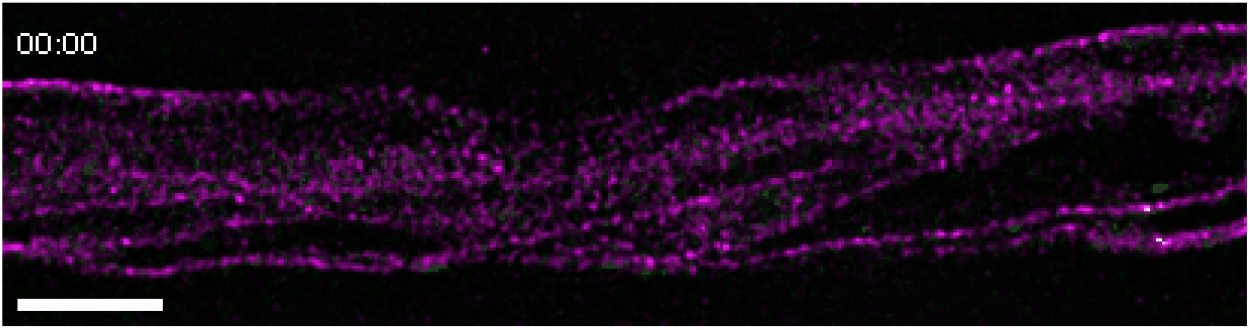
Timelapse of EB1b-mEGFP and mRuby2-TUB6 signal in the distal shank region. Images were captured every 10 seconds. Scale bar = 5 μm.

